# Urban birds become less fearful following COVID-19 reopenings

**DOI:** 10.1101/2023.01.04.522762

**Authors:** Eleanor S. Diamant, Ian MacGregor-Fors, Daniel T. Blumstein, Pamela J. Yeh

## Abstract

Following the COVID-19 pandemic, many people around the world stayed home, drastically altering human activity in cities. This exceptional moment provided researchers the opportunity to test how urban animals respond to human disturbance, in some cases testing fundamental questions on the mechanistic impact of urban behaviors on animal behavior. However, at the end of this “anthropause,” human activity returned to cities. How might each of these strong shifts affect wildlife in the short and long term? We focused on fear response, a trait essential to tolerating urban life. We measured flight initiation distance—at both individual and population-levels—for an urban bird before, during, and after the anthropause to examine if birds experienced longer-term changes after a year of lowered human presence. Dark-eyed juncos did not change fear levels during the anthropause, but they became drastically less fearful afterwards. These surprising and counter-intuitive findings, made possible by following the behavior of individuals over time, has led to a novel understanding that fear response can be driven by plasticity, yet not habituation-like processes. The pandemic-caused changes in human activity have shown that there is great complexity in how humans modify a behavioral trait fundamental to urban tolerance in animals.

## Main Text

In 2020, many countries in the world went into “lockdown” in response to the COVID-19 pandemic. With human mobility suddenly halted, these lockdowns drastically changed the dynamics of our cities and caused what has been coined as the “anthropause”^1^. While devastating for human communities, the absence of humans from the landscape provided a unique opportunity to study how animals respond to human activity, from the level of individual behavior to population dynamics to community composition. Likely because of the direct and indirect effects of human activity, such as vehicular traffic, collisions, light pollution, and noise pollution, some wildlife—specifically urban wildlife—adjusted their behaviors and patterns across the globe^1–6^. For example, during the initial pandemic lockdowns, urban white-crowned sparrows (*Zonotrichia leucophrys*) rapidly responded to the reduction in traffic noise by notably changing their songs to more high performing songs that are otherwise interrupted by urban noise^2^. Lockdowns have sporadically ended and re-occurred in different parts of the world, though human activity has broadly bounced back to pre-pandemic levels. As a result, urban animals are now faced with increased human activity and stressors following a long absence. By assessing their individual and population-level behaviors before, during, and after the anthropause, we can begin to understand how animals respond to dynamic human processes and stressors. We can also determine if and how this exceptional event continues to impact wildlife even after humans have returned to the landscape.

Determining how animals cope with urban stressors is essential to predicting wildlife response in the face of strong anthropogenic change^7,8^. Urbanization is a leading cause of habitat loss and biodiversity loss, though some animals manage to survive, adapt, and ultimately thrive in cities^9–12^. Though the underlying causes and associations for urban success vary across species and populations, tolerating humans is essential to urban life^13–15^. Indeed, at the population and species levels, we see that urban animals typically have a reduced fear of humans^16^. The mechanisms underlying this behavioral shift are challenging to parse out: in some organisms, this is due to habitat selection wherein individuals that are less fearful choose urban habitats with increased human activity and stress^17^. However, habituation-like processes that may underly within-generational behavioral plasticity—an individual’s propensity to shift their behavior in response to differences in their environments—can also explain this observed phenomenon^18^. Here, individuals exposed to human activity might decrease their fear response with increased exposure. Once in the city, the urban environment might select for individuals that are less fearful. Plasticity itself might be under selection if certain individuals express less fear upon exposure than others and may evolve if this has reproductive consequences^19,20^.

COVID-19 lockdowns and reopenings provided the opportunity for us to study the complex nature of how fear is affected by human activity, allowing us to test how plastic the response is in a successful urban bird: the dark-eyed junco (*Junco hyemalis*). This songbird, native to North America, began breeding in urban habitats in the past 20-40 years in Southern Californian cities^21–23^, likely independently. Across tested Southern California populations, they have a consistent decrease of fear response in comparison to non-urban conspecifics (Supplementary Information; Extended Data figure 1). By assessing fear response across lockdown conditions, we can then determine if and how non-evolutionary mechanisms, such as habituation-like processes that might lead to tolerance and habitat selection whereby tolerant individuals settle around humans while less tolerant ones avoid humans, as well as evolutionary processes like selection on plasticity itself, impact fear. Further, the relatively sudden reintroduction of humans to the landscape provided us with the opportunity to assess if dark-eyed junco behavior returned to a pre-pandemic “normal” or if lockdowns shifted how this urban bird behaves and copes with human presence long-term.

We tested individual and population-level fear response in urban dark-eyed juncos before, during, and after COVID-19 closures to understand the immediate and longer-term effects of COVID-19 on other animals and to test fundamental questions in urban behavioral adaptation. If lower urban fear response is due to habituation, we expected fear response to lower during the COVID-19 closures and increase following reopening. On the other hand, if lower urban fear response is due to habitat selection and is less plastic, we expected fear response to remain unchanged with respect to the closures. If both habitat selection and habituation play a role, as might be seen if birds that have relatively lower fear response and a plastic habituative response are selected for, we expected fear response to increase during the closures, though not to the level of non-urban birds, and to decrease after reopening at the individual and population level. None of these hypotheses were supported by our findings. Instead, we found that, at the population level and individual level, urban birds did not change their behavior during the COVID-19 closures but became significantly less fearful of humans following reopenings in comparison to pre-pandemic baselines.

Because we have a longitudinal study site in urban Los Angeles, dark-eyed juncos at the University of California Los Angeles (“UCLA”) were individually distinguishable by unique colored leg bands. In March 2020, UCLA closed classes and research, except for essential researchers. UCLA remained remote until Fall 2021, when classes resumed on campus. Campus closures caused human activity to be approximately 7x lower than “normal” in 2021^24^, and therefore even lower during 2020 at the height of Los Angeles lockdowns and UCLA campus restrictions. We assessed the fear levels of individually identifiable birds using a flight initiation distance assay^25,26^ repeatedly during the 2018-2022 breeding seasons (January-June/July). These breeding seasons encompassed pre-pandemic, city lockdowns starting in March 2020, campus closures continuing through 2021, and post-reopening times in 2022. We categorized these time periods as “pre-pandemic” (2018-March 14, 2020), “2020 anthropause” (March 15, 2020-July 2020), “2021 anthropause,” and “post-anthropause.”

## Results

At the population-level, fear response remained relatively unchanged during campus closure in comparison to pre-pandemic levels (*N*=404). Upon campus reopening in the 2021-22 academic year, fear response in the 2022 breeding season was significantly reduced compared to measurements from prior breeding seasons (Fig. 1). We used a regression tree method to determine relevant variables explaining fear response and subsequently fitted a generalized linear mixed model (GLMM) (see Methods). Our fixed effects included campus closure status (2018-2020 pre-pandemic, 2020 anthropause, 2021 anthropause, 2022 post-anthropause), relative change in human activity in Los Angeles (average change across residential and non-residential activity from a pre-lockdown baseline quantified using Google COVID-19 Community Mobility Reports^27^), the trial number of the focal bird, and starting distance. Individual bird ID was a random effect (Supplementary Table 1). We found that campus closure status (*X*^2^=20.4, df=3, p=0.0001) strongly and significantly affected fear, but relative change in human activity (*X*^2^=1.48, df=1, p=0.22), trial number (*X*^2^=2.12, df=1, p=0.15) and starting distance (*X*^2^=0.03, df=1, p=0.85) did not. Additionally, there was significant individual variation in fear (sd=0.23, p<0.0001; Extended Data figure 2). Collectively, individual juncos varied in their fear response, but the reopening and increase of human activity following restrictions significantly lowered fear across the population (p<0.001 in 2020 anthropause and 2021 anthropause fearfulness compared with post-anthropause fearfulness, and p=0.008 in pre-pandemic fearfulness compared with post-anthropause fearfulness; Fig. 1; Extended Data figure 2; Supplementary Table 1).

**Fig. 1.**
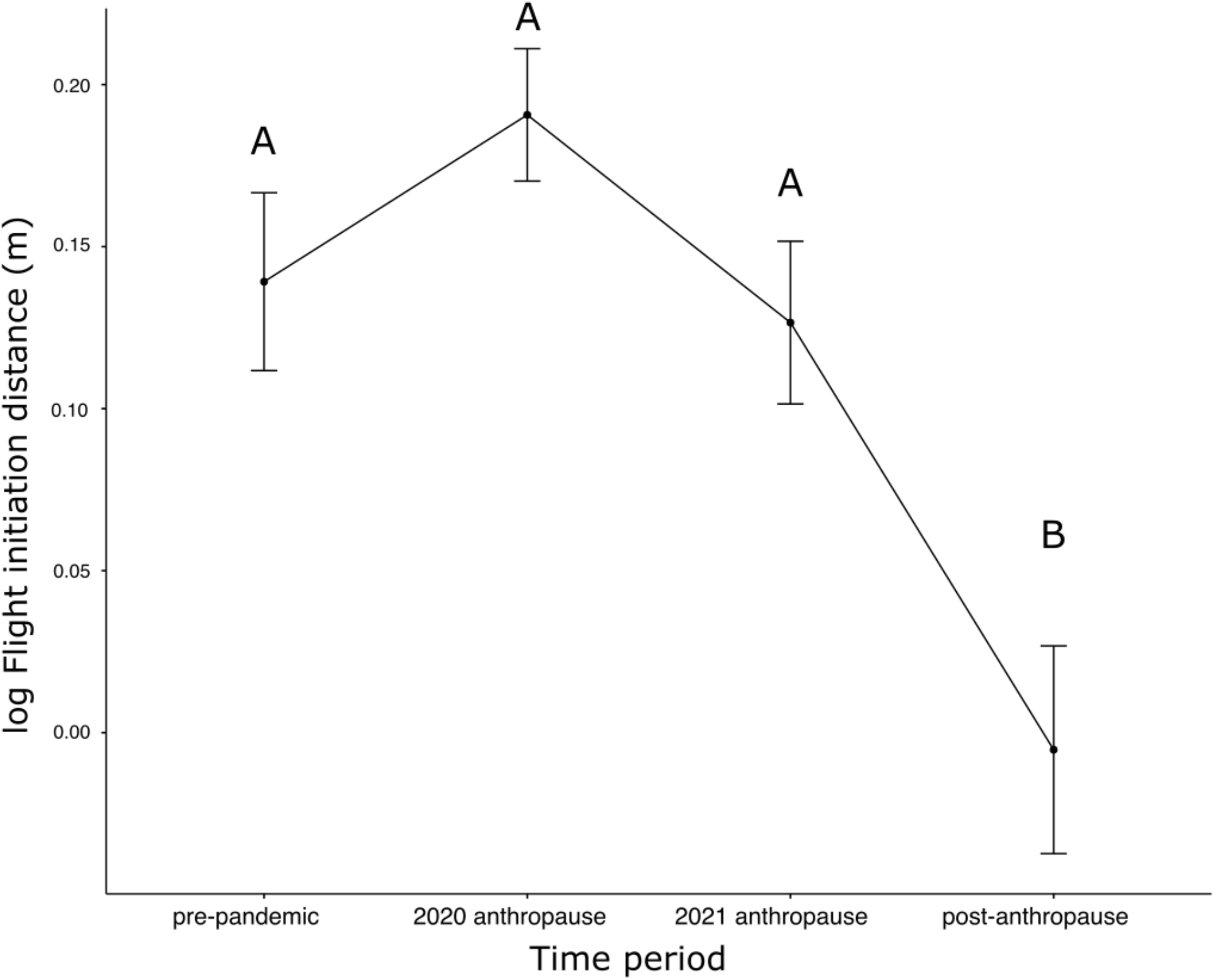
Population-level fearfulness remained unchanged during lockdowns but decreased following reopenings. Population-level flight initiation distance (m) before (*n*=71), during (*n*_2020_=135; *n*_2021_=131), and after (*n*=67) the anthropause. The dark-eyed junco population at University of California Los Angeles (UCLA) did not shift their FID across the anthropause (GLMM contrast: p>0.05). FID significantly dropped in the 2022 post-anthropause environment in comparison to both years in the anthropause and the pre-pandemic baseline (GLMM contrasts: p<0.0001 for pre-pandemic, 2020 anthropause, and 2021 anthropause compared to 2022 post-anthropause). Flight initiation distance (FID) data are log_10_ transformed for visual aid, but not in the formal statistical analysis. Data points represent mean log_10_ FID ± standard error for each time period assessed. Groups with the same letter are not statistically significantly different from each other. Groups with different letters are statistically significantly different.

The patterns seen at the population level were repeated at the individual level (Fig. 2; Supplementary Table 2). A subset of individuals was tested repeatedly before, during, and after pandemic closures. While individuals varied in their fear response once lockdowns occurred, individuals nearly universally became less fearful following campus reopening. These results are consistent with the hypothesis that behavioral plasticity in fear response explained the pattern at the population-level. While we expected individual birds to become more tolerant to humans during the closures and return to a pre-pandemic fear response following reopening, we found that urban dark-eyed juncos had a surprisingly plastic response to increased human activity, but not to decreased human activity.

**Fig. 2.**
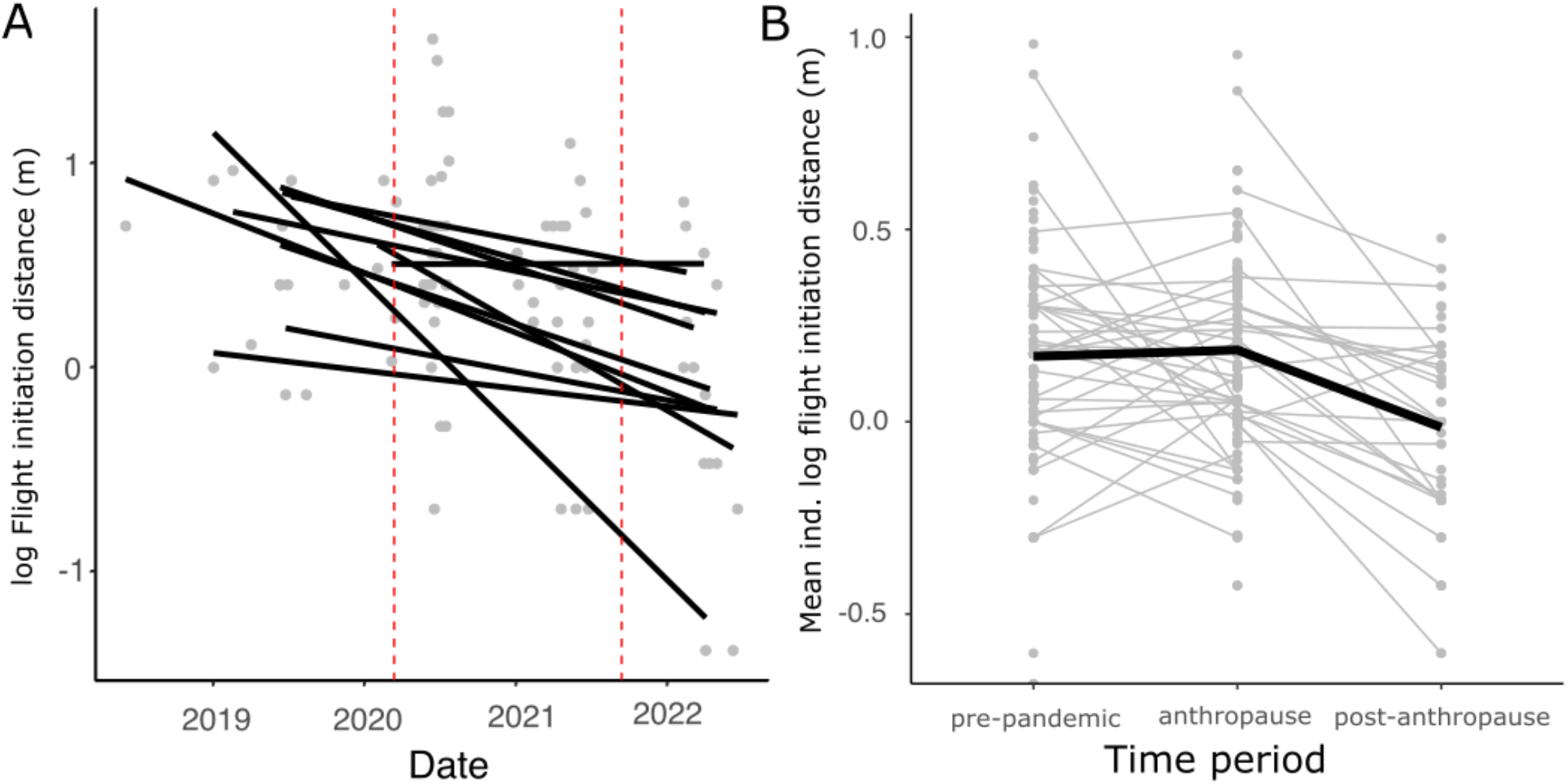
Individuals became less fearful after reopenings in comparison to before the pandemic closures. Individual shifts in flight initiation distance (FID) before, during, and after anthropause. A. FID measurements only for individuals that were tested repeatably for at least one time point in each period: before, during, and after the campus closures (*n*=11). Trends reflect a similar decrease in FID from before the pandemic to the end of the anthropause. Lines are fitted linearly to demonstrate the change from before to after the anthropause. Each line represents one individual. Dashed red vertical lines denote the beginning and end of the anthropause, respectively. A GLMM, only including birds tested across all time periods and accounting for potential habituation (by including trial number), and treating the anthropause as a single category were similar. Here, differences in pre-pandemic fearfulness compared with post-anthropause fearfulness were marginally significant (p=0.09) and differences in anthropause compared to post-anthropause fearfulness were significant (p=0.02) B. Mean FID values per individual (gray) in pre-pandemic, anthropause, and following reopening—”post-anthropause”— time periods. The thick black line represents shifts across all individuals sampled repeatedly between the pre-pandemic and anthropause (*n*=33) or the anthropause and post-anthropause time periods (*n*=24). These demonstrate pairwise shifts in mean fear response for each individual to account for individuals that might not have been tested in one of the time periods. A GLMM accounting for potential habituation (by including trial number) and considering the anthropause as a single category, revealed significant pairwise differences between pre-pandemic and post-anthropause environments (p=0.008) and between the anthropause and post-anthropause environments (p<0.001). Though FIDs were not log_10_ transformed in our analyses (given their gamma distribution), they are log_10_ transformed here for visual ease.

## Discussion

We found that fear response did not shift at the height of the COVID-19 closures, in comparison to before the pandemic, but that the re-introduction of humans led to a decrease in fear both at the population-level and at the individual-level. Thus, this urban bird did not become more like its wildland counterparts without human presence but rather, when faced with more human activity, became even less fearful than pre-pandemic levels, which is already less fearful than wildland birds. That dark-eyed juncos did not increase fearfulness with decreased human activity, even amongst birds who hatched during the COVID-19 closures (Supplementary Information; Extended Data figure 3), suggests that lower urban fear response is not dynamically driven by habituation-like processes—if these processes have a role in explaining tolerance at all. Indeed, habituation-like processes were controlled for at the individual level in all our analyses and do not explain the population-wide pattern we found. Rather, fear response expression is likely more complex and suggests that studying their ontogeny will be particularly illuminating. It could be that urban colonists from non-urban origins might have a lower baseline fear response in comparison to the non-urban population at large, and potentially decrease their fear response with human activity. We note, that in contrast to virtually all other studies of anthropause effects in birds (e.g.,^1,3,4,28–30^) that used unmarked birds and were unable to focus on individuals, these insights emerged only from a detailed, longitudinal study of individuals.

A population increase coupled with a re-introduced landscape of fear might have led to higher competition for resources in 2022 and thus trade-offs favoring increased foraging despite higher perceived predator (human) risk. Changing patterns of human mobility shifted birds’ use of space broadly across lockdowns^5^, suggesting that human presence affects the habitability of urban spaces. Entering into COVID-19 lockdowns and reopenings, contrasting shifts in human activity—one increasing human activity and one decreasing human activity—led to contrasting fitness consequences reflecting this shift in the landscape of fear: great tits (*Parus major*) in an area with lower human activity had higher reproductive output than that with higher human activity^31^. However, there was no evidence for increased fitness by means of increased nestling condition and nest success when comparing 2021 and pre-pandemic 2019 breeding seasons in this population^24^. Additionally, there were no changes in aggressive interactions in the population following reopenings in 2022 and pre-pandemic 2019 breeding seasons^24^, suggesting that fear response is not a by-product of shifting behavioral strategies due to a different socioecological context or due to indirect effects on predator density during the anthropause. Urban song rapidly shifted during San Francisco’s lockdowns in a related species (white-crowned sparrows) potentially because there was a clear communicative signal being interrupted by urban stressors^2^. The relationships that exist between human activity and other urban behaviors appear more nuanced.

Fear responses could vary because ecological conditions changed, altering the trade-offs in escape behavior following reopenings. Recent drought conditions in Southern California might have made urban birds more reliant on anthropogenic food to buffer declines in natural food resources—as was the case in an urban monkey^32^—leading to a higher tolerance of human presence in 2022. However, urban areas act as a buffer to arid conditions because of irrigation, supporting larger populations and diversity of arthropods^33^. Additionally, UCLA is an irrigated and green campus in an affluent area, which in turn is associated with increased irrigation and higher plant and bird diversity relative to non-urban arid conditions^34^. Thus, drought conditions in 2022 might not have caused strong detrimental effects, if any, to local urban resource distribution. Emergency regulations in California limiting turf irrigation only began in June 2022, making this a particularly unlikely explanation for lower fear response in the time period we were sampling, but something that could be accounted for in future studies given recent water-use restrictions.

Alternately, we could have been measuring the level juncos were distracted by stimuli. Indeed, escape behavior can vary based on the number of stimuli as prey must divide their attention. With high human density and disturbances, prey can become distracted and either fail to respond as rapidly to an approaching threat or flee more rapidly^35,36^. Here, fear *response* in juncos would reflect the focal bird being more or less distracted, rather than more or less fearful. If distraction by the sudden increase in humans was responsible for the reduction in FID, we might expect that distracted animals were unable to detect an approaching human and therefore tolerated closer approach. Nonetheless, if fear response were solely modulated by distraction, we would have expected a higher fear response during the anthropause than after reopening; this pattern was not seen. Thus, juncos behaved surprisingly differently following the re-introduction of high human activity, though it could be due to them filtering stimuli differently than before the pandemic.

We propose two novel hypotheses explaining how fear response develops and is modulated in urban-adapted birds: urban bird fear response is either a ratchet or a spring. Birds that hatched during the anthropause mirrored the population as a whole: they became less fearful with increasing human activity following reopenings rather than expressing fearfulness at the same level as second year birds did before the pandemic, despite differences in their early life environment. Similarly, the population as a whole became less fearful than their pre-pandemic fear levels following reopenings (Supplementary Information). Thus, the prolonged absence of human activity followed by a rapid increase, rather than recurrent exposures to human activity, could be driving the expression of this fear response.

If a fear response acts as a spring that returns to pre-existing baseline with continuous exposure, we would expect that dark-eyed juncos will eventually re-sensitize to human activity and return to a pre-pandemic intermediate baseline. Alternatively, fear responses could change like a ratchet where each burst of rapid increases in human activity an urban bird is exposed to could lead to lower fear response. Rather than dynamically returning to a pre-pandemic baseline, a long absence coupled with a rapid burst of human activity drives an increase in human tolerance. Testing these hypotheses requires on-going study.

Collectively, our results suggest that changes in fear responses might not be as predictable as we might expect, and likely depends on *which* individuals and *how* their behaviors develop and shift in combination with strong and rapidly shifting collective human behaviors. Only through studies on individual animals tracked over time can we understand the mechanisms underlying population response, which cannot be confirmed from contradictory broadscale patterns found in metanalyses^28,30^. While the anthropause created much human hardship, it offered a unique opportunity to identify an important new avenue of ontogenetic research that can create insights which will help us better conserve biodiversity in a rapidly changing, human-dominated world.

## Online Methods

### Study Sites

We conducted field work at urban and non-urban sites in Southern California between 2017 and 2022. We assessed birds across three metropolitan areas: Santa Barbara County (“Santa Barbara”), Los Angeles County (“Los Angeles”), and San Diego County (“San Diego”) (Extended Data figure 2). At each of these metropolitan areas, we sampled birds at the local University of California (University of California Santa Barbara (UCSB), Los Angeles (UCLA), and San Diego (UCSD), respectively). In Los Angeles, specifically, we also sampled birds across the city in Occidental College and parks of various sizes across the urban core. Dark-eyed juncos likely began breeding in San Diego in the early to mid 1980s, in Los Angeles in the early to mid 2000s, and in Santa Barbara in the early 2010s. These sites were compared to non-urban, mountainous sites that are indicative of their historic breeding range ^23,37^: the UC Stunt Ranch Reserve in the Santa Monica Mountains, the UC James San Jacinto Mountain Reserve, and the Angeles Forest in the San Gabriel Mountains.

### Individual Color Banding

We captured and banded local dark-eyed juncos at each site at the start of territorial singing – around mid-January in urban sites and April in non-urban sites – to July 2017. Birds were captured between 6:30 and 11:00. They were lured into mist-nets using playback of junco song recorded at UCLA in 2018 or from the MacCauley Library (Cornell University). Each junco was fitted with 3 color bands and 1 aluminum USGS band in a unique combination. Birds were aged by molt limits as “second year,” “after second year,” or “after hatch year” (when age could not be determined) and sexed by cloacal protuberance or brood patch. When birds were not in breeding condition, they were sexed by plumage, which was later confirmed by behavior (singing or exhibiting nesting behavior). All birds were released after processing.

### Flight Initiation Distance Assays

We determined flight initiation distances (FIDs) for each bird following^25,26^. All FID tests were conducted by ESD. A marker was dropped at first site of the focal bird. The researcher walked at a steady pace of ~0.5m/sec towards the bird. A second marker was dropped at the point the researcher was when the bird flew or hopped away, and then a third where the bird was when it flew or hopped. We recorded the starting distance (the distance between the first marker and the third), FID (the distance between the second and third). Effort was made to universally assay juncos in instances with low human activity (<15 humans walking in the vicinity while assays were conducted, except for 18 data points in areas rarely empty during the academic year) and with no other humans or juncos between the investigator and the focal bird. Trial number per individual was determined for each fear response assay and varied between 1 to 10 trials per individual.

We conducted FID tests in 2018 and 2019 in non-urban sites. At sites other than UCLA, we conducted FIDs before COVID-19 lockdowns in 2018, 2019, and up to March 2020, as the campuses closed following “safer-at-home” measures. At UCLA, we conducted FIDs on birds in 2018, 2019, 2020 (*n*_pre-pandemic_=71; *n*_2020 anthropause_=135), 2021 (*n*=131), and 2022 (*n*=67). Most FIDs were conducted during the pre-breeding and breeding seasons—between January and July—as birds were more conspicuous, easier to find, and not in wintering flocks, which has been found to affect FIDs in other songbirds ^38^. We measured FIDs of juncos before COVID-19 restrictions started on 14 March 2020 (i.e., “pre-pandemic”). In-person classes were canceled on that date, and the campus was mostly closed thereafter (i.e., “during anthropause”). During this time, we attempted to re-measure FIDs for individuals every 2 weeks, though this was not possible universally due to the spontaneous nature of FID testing. We re-assessed individuals following COVID-19 lockdowns from January to June 2022 following UCLA returning to in-person instruction (i.e., “after anthropause”). We categorized these time periods as “pre-pandemic” (2018-March 14, 2020), “2020 anthropause” (March 25, 2020-July 2020), “2021 anthropause,” and “post-anthropause” (2022).

Because we were also tracking reproductive behavior prior to COVID-19 lockdowns, we know with certainty that at UCLA all chicks were hatched following the cancelation of in-person classes, and thus a significant decrease in human activity. As such, all birds banded at UCLA in 2021 and 2022 that were in their second-year plumage were likely to have been hatched and fledged during the COVID-19 lockdown, with minimal human activity relative to older birds on campus. We also followed reproductive activities in 2021 and banded chicks in the nest, some of which returned to campus and were assayed as second-year birds.

### Measuring Human Activity

We conducted human pedestrian surveys at UCLA to confirm that human activity was lower during campus closures than when classes are in session and in person. We surveyed 12 points across campus twice per week, once in the morning and once in the afternoon during the anthropause (May-July 2021) and the campus was “back to normal” (March-July 2022). Each survey lasted two minutes and all individuals, vehicles, and dogs crossing the observer’s eyeline were counted.

To gauge human activity shifts in the city as a whole, we accessed Google’s Community Mobility Reports^27^. This dataset compiled data from smartphones to determine where users spent time across different place categories compares aggregated data to a pre-pandemic baseline for a given community (in this case, Los Angeles) per day. We used the categories: mean change in activity (averaged across categories) and time spent in residential places as variables in our analysis. Because this dataset only began in February 2020, we calculated the mean and standard deviation for pre-pandemic levels (15 February to 14 March 2020) and randomly generated a normal distribution with the calculated mean and standard deviation. We imputed Los Angeles Community Mobility data for each date before 15 February 2020 randomly from this normal distribution.

### Statistical Analysis

To test if urban dark-eyed juncos across Southern Californian cities have parallel shifts in FID in comparison to non-urban dark-eyed juncos, we fitted a generalized linear mixed model (GLMM) using a gamma distribution and an inverse link function. We included city and starting distance as fixed effects and bird ID as a random effect. For this analysis, we only included dark-eyed juncos in Los Angeles (*n*=119), San Diego (*n*=33), and Santa Barbara (*n*=13) that were assayed before the COVID-19 pandemic. We aggregated data (*n*=25) across non-urban populations to compare to. We then determined within model contrasts to test if urban populations were different from each other and from non-urban juncos.

To determine which variables were important to include in GLMMs, we first ran a regression tree analysis. Here, our dependent variable was flight initiation distance (m). Independent variables we included were: sex, month, UCLA’s anthropause status+year (2018-2020 “pre-pandemic,” “2020 anthropause,” “2021 anthropause,” 2022 “post-anthropause”), Google’s mean change in activity in Los Angeles, Google’s time spent in residential places in Los Angeles, and starting distance. Based on this analysis, UCLA’s anthropause status+year, mean change in community mobility, and starting distance were variables that were found to diagnose differences in flight initiation distance. Mean change in activity and time spent in residential places were associated with each other, because mean change in activity is calculated using time spent in residential places, in combination with other variables. The regression tree analysis found that mean change in activity was a more important variable than time spent in residential areas. Thus, we included mean change in activity in our GLMM.

We then fitted a GLMM with a gamma distribution and an inverse link, fitting the distribution of our data. We included UCLA’s anthropause status+year, a given individuals’ trial number, mean change in community mobility use with the Los Angeles Google Community Mobility Reports data, and starting distance as fixed effects and bird identity as a random effect. We visualized individual shifts by subsetting juncos who were assessed before and during the anthropause and/or during and after the anthropause to determine if trends at the population level were repeated at the individual level (fig. 2). Due to lower sample sizes for samples within individuals across time periods, we combined 2020 and 2021 anthropause categories into one category. To determine if patterns in these data were driven by habituation-like processes, we fitted the same GLMM, but only considered 3 anthropause categories (i.e., “before,” “during,” and “after” the anthropause) instead of differentiating between 2020 and 2021. All other variables remained the same.

To assess early-life effects, we compared four cohorts of second-year juncos’ FID at UCLA. Second-year birds assayed in 2019 (*n*=15) hatched and were assayed in a high-human-activity environment. Second-year birds assayed in 2020 (*n*=34) hatched with high human activity and were assayed in an anthropause environment. Second-year birds assayed in 2021 (*n*=11) hatched and were assayed in an anthropause environment. Second-year birds assayed in 2022 (*n*=10) hatched in an anthropause environment yet were assayed in a high-human-activity environment. We fitted a GLMM with a gamma distribution and an inverse link function. The fixed effects were “year second-year bird was assayed” (pre-pandemic, 2020 anthropause, 2021 anthropause, 2022 post-anthropause) and starting distance. Bird ID was included as a random effect. We calculated within-model contrasts to determine if there were significant differences in FID between groups. We tested the assumptions of all models by testing for normality and linearity of residuals, as well checking for multicollinearity between independent variables. Assumptions were met for all models.

All tests were done in R v. 4.2.2^39^. Regression trees were built and tested using R packages RSAMPLE^40^ and RPART^41^. GLMMs were built and analyzed using LME4^43^. Within-model contrasts were calculated using R packages GMODELS^44^ and MULTCOMP^45^.

## Data and materials availability

All data and code are available through Figshare and will be publicly accessible upon acceptance.

## End Notes

## Acknowledgments

We are thankful to Samuel Bressler, Marlene Walters, Taylor Kang, and Wilmer Amaya-Mejia for their participation in the banding effort. We also thank all undergraduate research assistants from 2018-2022 who have helped with banding stations, particularly Christina Cen, Carissa Reulbach, and Julia Lung. We thank the American Ornithological Society Hesse Award, University of California Los Angeles Stunt Ranch and the La Kretz Center Research Award, Pasadena Audubon Society, Santa Monica Audubon Society, and The Western Section of the Wildlife Society for funding.

## Author contributions

ESD contributed conceptualization, methodology, investigation both in the field and data analysis, visualizing, funding acquisition, writing the original draft, and subsequent editing. IM-G contributed data analysis, visualization, and manuscript editing. DTB contributed methodology and manuscript editing. PJY contributed conceptualization, methodology, visualization, funding acquisition, supervision, and manuscript editing.

## Competing interests

Authors declare that they have no competing interests.

Supplementary Information is available for this paper.

Correspondence and requests for materials should be addressed to Eleanor Diamant.

