## Supplementary Material for "Urban birds become less fearful following COVID-19 reopenings"

### Supplementary Text

#### Fear response across Southern Californian populations

Urban populations across Southern California exhibited lower, yet similar FID to each other in a pre-pandemic baseline, in comparison to non-urban dark eyed juncos aggregated across non-urban populations (Extended Data figure 1). Dark-eyed junco population was a significant effect in the model ($X^2$=41.8, df=3, p<0.0001) and starting distance was not ($X^2$=2.59, df=1, p=0.108). All urban populations had non-significant differences in FID (p>0.1), but each was significantly lower than non-urban juncos (p<0.01), suggesting that time since urban colonization and population does not affect an urban junco population from shifting fear response lower.

#### Fear response across age cohorts

FID significantly varied by anthropause stage+year when assessing second-year bird fear response ($X^2$=8.10 df=3, p=0.03; starting distance: $X^2$=3.67; df=1, p=0.06; trial number: $X^2$=1.84; df=1, p=0.17). 2022 second-year juncos—those that hatched during the anthropause yet had exposure to human activity in adulthood—had a substantially lower FID in comparison to all other groups. These results were marginally significant in pairwise contrasts (pre-pandemic – post-anthropause: p=0.02; 2020 anthropause – 2022 post-anthropause: p=0.04; 2021 anthropause – 2022 post-anthropasue: p=0.09; Extended Data figure 3). However, all other groups did not significantly differ from each other ($p>0.90$). Low sample sizes likely affected power to detect differences between groups using an $\alpha=0.05$.

### Supplementary Tables

**Supplementary Table 1.** Generalized linear mixed model (GLMM) results for population-level differences in flight initiation distance (FID) across years and anthropause status (before, during, after).

| ***Fixed effects*** | $X^2$ | *df* | *p-value* |
|---|---|---|---|
| Mean change in Los Angeles mobility across measured mobility categories | 1.48 | 1 | 0.22 |
| Trial number | 2.12 | 1 | 0.15 |
| Anthropause+Year | 20.42 | 3 | **0.0001** |
| Starting distance | 0.03 | 1 | 0.85 |

| ***Random effects*** | *variance* | *sd* | *p-value* |
|---|---|---|---|
| Bird ID | 0.05 | 0.23 | **< 0.0001** |
| Residual | 0.24 | 0.49 | |

| ***Number of observations*** | 402 |
|---|---|
| ***Log-likelihood*** | -436.6 |

| ***Pairwise contrasts: Anthropause+Year*** | *estimate* | *se* | *z-score* | *p-value* |
|---|---|---|---|---|
| pre-pandemic – 2020 anthropause | -0.09 | 0.09 | -0.71 | 0.65 |
| pre-pandemic –2021 anthropause | -0.10 | 0.10 | -1.32 | 0.10 |
| pre-pandemic –post-anthropause | -0.36 | 0.11 | -3.13 | **0.008** |
| 2020 anthropause – 2021 anthropause | -0.06 | 0.05 | -1.35 | 0.12 |
| 2020 anthropause – post-anthropause | -0.29 | 0.07 | -4.42 | **< 0.001** |
| 2021 anthropause – post-anthropause | -0.23 | 0.06 | -3.80 | **< 0.001** |

GLMM is fitted to a gamma distribution with an inverse link. P-values for random effects were calculated by comparing models with and without the given random effect using ANOVA. Pairwise contrasts for anthropause stage categories for the model are included. Significant p-values are in bold. Significant p-values for pairwise contrasts reflect significant differences in FID within the model between the groups contrasted.

**Supplementary Table 2.** Generalized linear mixed model (GLMM) results for individual-level differences in flight initiation distance (FID) across years and anthropause status (before, during, after) with 3 categories for COVID-19 closure status.

| ***Fixed effects*** | $X^2$ | *df* | *p-value* |
|---|---|---|---|
| Mean change in Los Angeles mobility across measured mobility categories | 1.73 | 1 | 0.19 |
| Trial number | 5.22 | 1 | **0.02** |
| Anthropause status | 18.6 | 3 | **< 0.0001** |
| Starting distance | 0.07 | 1 | 0.79 |

| ***Random effects*** | *variance* | *sd* | *p-value* |
|---|---|---|---|
| Bird ID | 0.05 | 0.22 | **< 0.0001** |
| Residual | 0.24 | 0.49 | |

| ***Number of observations*** | 402 |
|---|---|
| ***Log-likelihood*** | -437.6 |

| ***Pairwise contrasts: Anthropause*** | *estimate* | *se* | *z-score* | *p-value* |
|---|---|---|---|---|
| pre-pandemic – anthropause | -0.08 | 0.09 | -0.88 | 0.63 |
| pre-pandemic –post-anthropause | -0.33 | 0.11 | -2.94 | **0.008** |
| anthropause – post-anthropause | -0.25 | 0.06 | -4.28 | **< 0.001** |

GLMM is fitted to a gamma distribution with an inverse link. P-values for random effects were calculated by comparing models with and without the given random effect using ANOVA. Pairwise contrasts for anthropause stage categories for the model are included. Significant p-values are in bold. Significant p-values for pairwise contrasts reflect significant differences in FID within the model between the groups contrasted.

**Extended Data Figures**

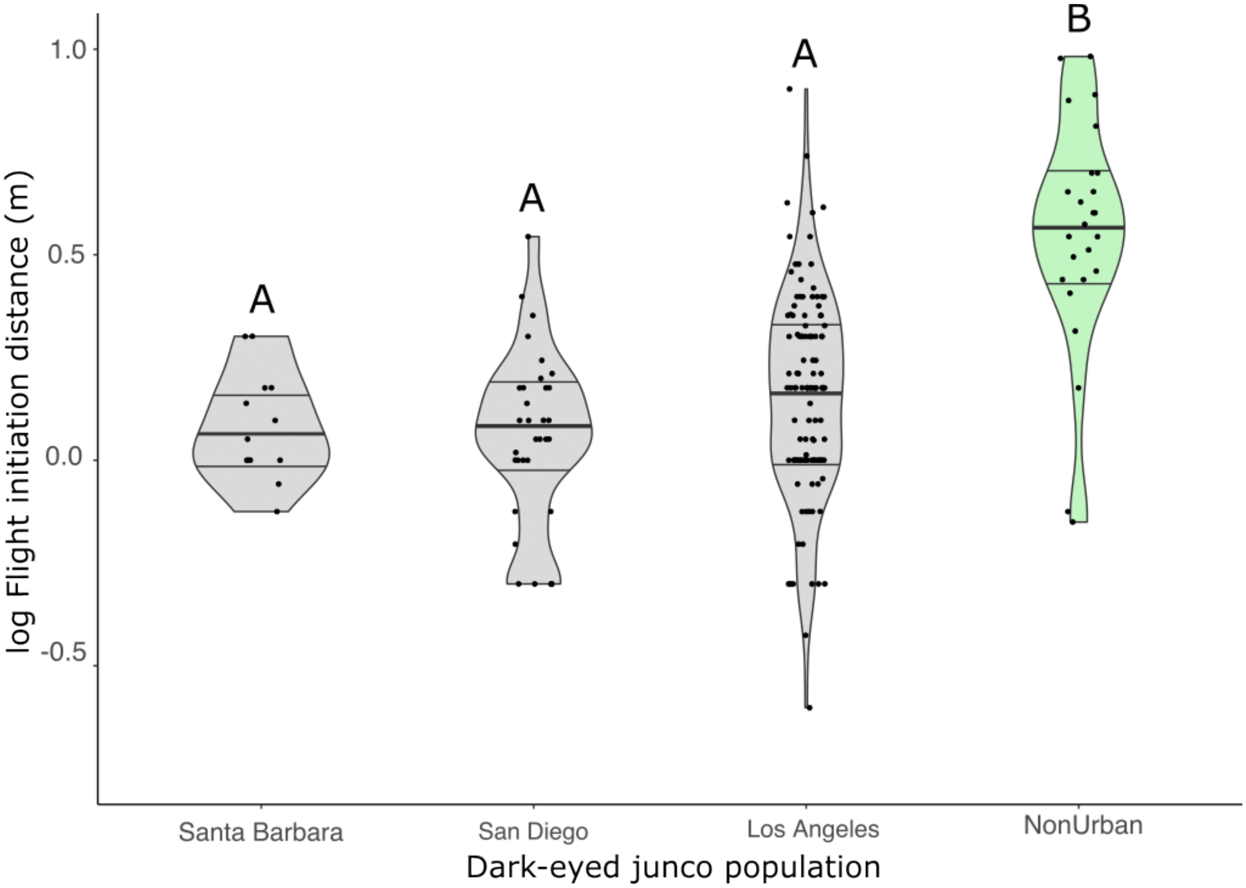


**Extended Data Fig. 1. Flight initiation distance (m) was lower across urban dark-eyed junco populations in comparison with non-urban populations in Southern California.** Flight initiation distance in Los Angeles (*n*=119), San Diego (*n*=33), and Santa Barbara (*n*=13) were all significantly lower than that of non-urban dark-eyed juncos (*n*=25) ($p<0.01$), yet not significantly lower from each other ($p>0.05$). Flight initiation distance data are log-transformed for visual aid, but not in the analysis. Grey represents urban populations and green represents non-urban populations. Shaded violin plots represent the data distribution. Lines within the violin plots represent 25%, 50% (thicker center line), and 75% quantiles. Data points per category are jittered. Groups with the same letter are not statistically significantly different from each other. Groups with different letters are statistically significantly different.

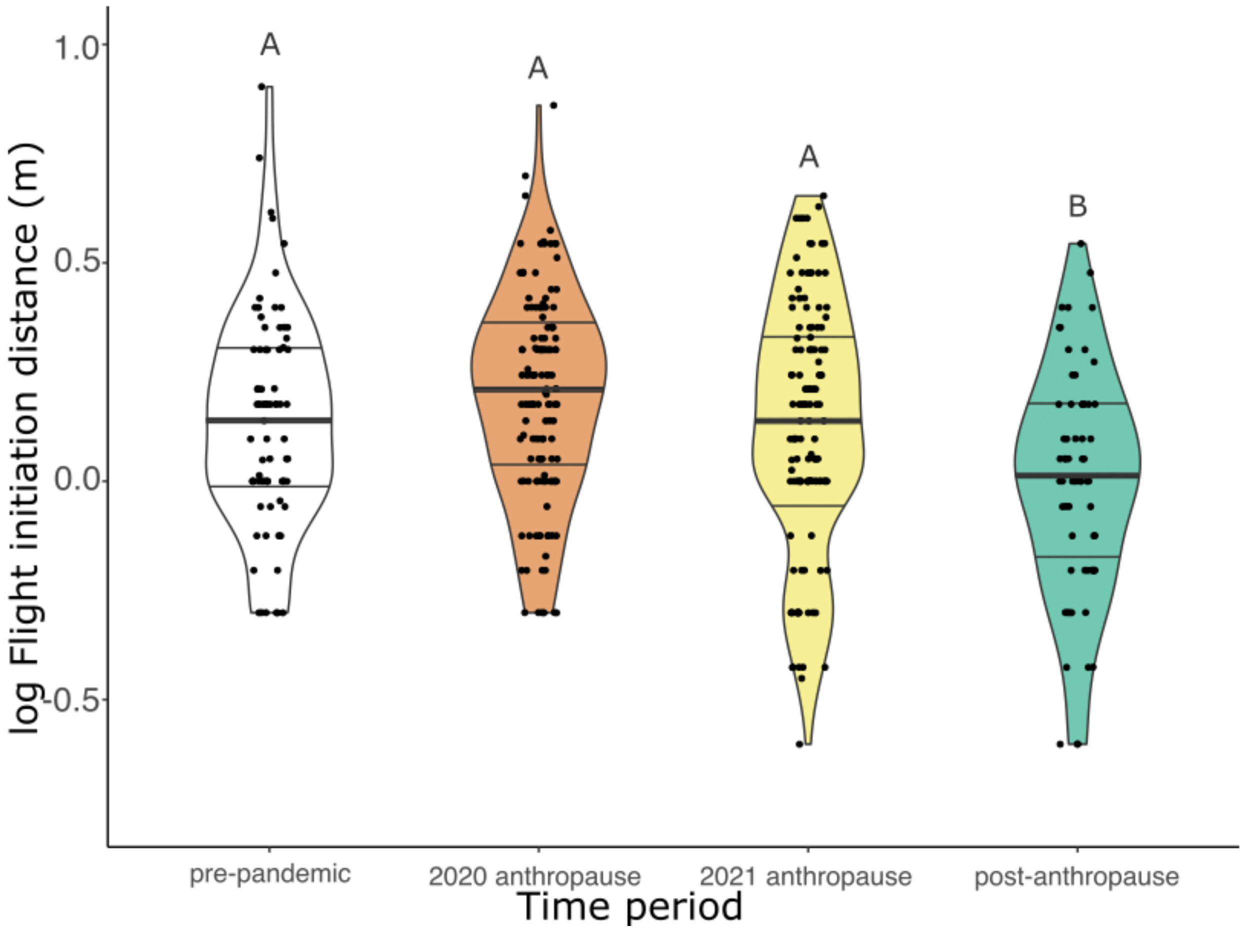


**Extended Data Fig. 2. Population-level fearfulness remained unchanged during lockdowns but decreased following reopenings, though exhibits strong variation across time.** Population-level flight initiation distance (m) before ($n_{2018+2019+2020\ \text{pre-pandemic}}$ =71), during ($n_{2020\ \text{anthropause}}$=135; $n_{2021\ \text{anthropause}}$=131), and after ($n_{2022}$=67) the anthropause. The dark-eyed junco population at University of California Los Angeles (UCLA) did not shift their FID across campus closures (GLMM contrasts: $p>0.05$. FID significantly dropped in the 2022 post-anthropause environment in comparison to both years in the anthropause and the pre-pandemic baseline (GLMM contrasts: $p<0.01$). Flight initiation distance (FID) data are log-transformed for visual aid, but not in the analysis. Shaded violin plots represent the data distribution, with white represented a pre-pandemic baseline, orange-red representing closures in UCLA and Los Angeles as whole, yellow representing the lifting of some but not all restrictions, and green representing full reopenings and the end of the anthropause. Lines within the violin plots represent 25%, 50% (thicker center line), and 75% quantiles. Data points per category are jittered. Groups with the same letter are not statistically significantly different from each other. Groups with different letters are statistically significantly different.

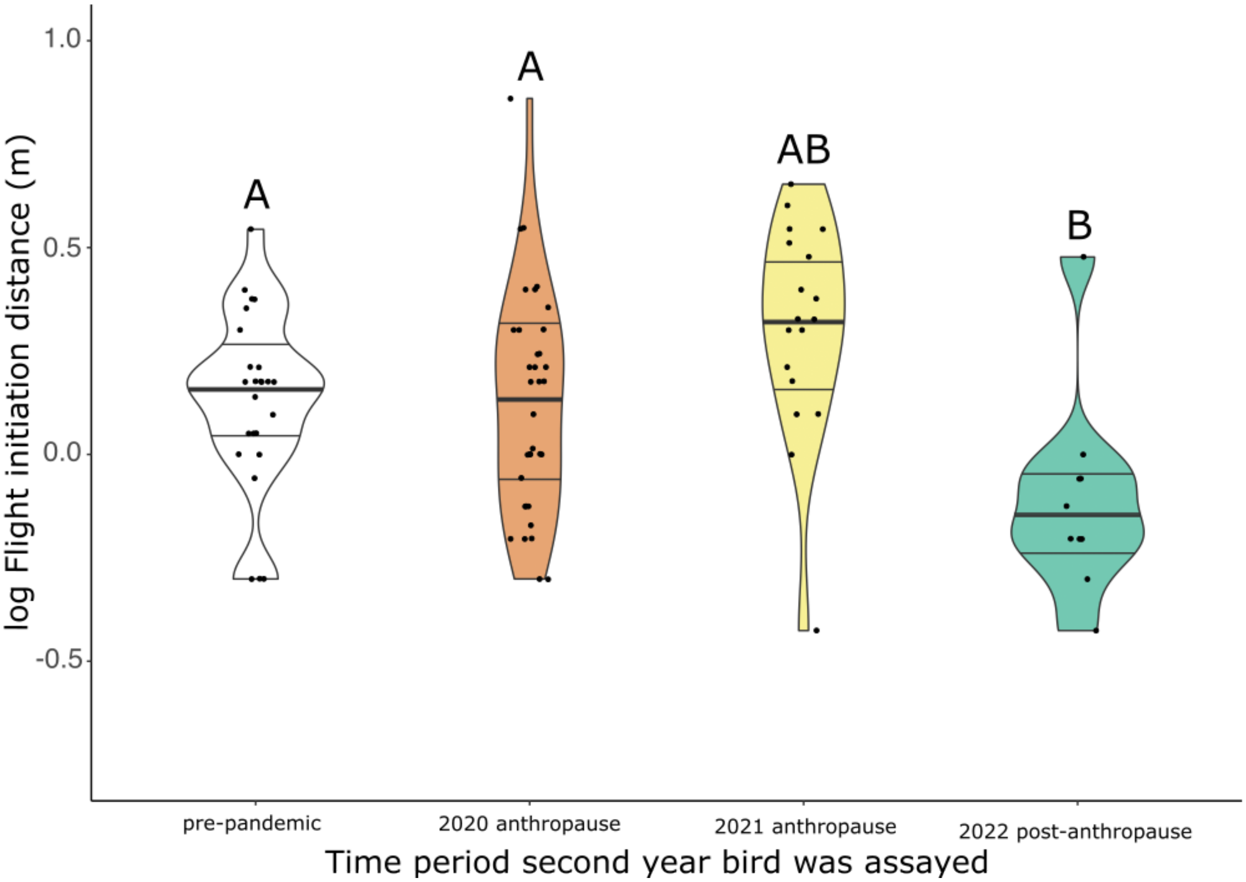


**Extended Data Fig. 3. Early life and adulthood exposure to human activity does not affect adult fear response.** Each column corresponds to only second-year birds assayed in different years in a natural full factorial experiment. The left column represents second-year birds assayed in 2019 (*n*=15) hatched and were tested in a pre-pandemic, high human activity environment. Second-year birds assayed in 2020 (*n*=34) hatched in an environment with high human activity yet were assayed as adults during the anthropause. Next, second-year birds assayed in 2021 (*n*=11) hatched and were tested during the anthropause, never having been exposed to high human activity. Finally, the right column represents second-year birds assayed in 2022 (*n*=10): hatched during the anthropause and naively exposed to high human activity in adulthood. 2022 second-year birds have a marginally significantly shorter FID than other groups (2019-2022: p=0.02; 2020-2022: p=0.04; 2021-2022: p=0.09), and all other groups were not significantly different from each other (p>0.9). Data are restricted to dark-eyed juncos at UCLA. Shaded violin plots represent the data distribution. Flight initiation distance data are log-transformed for visual aid, but not in the analysis. Shaded violin plots represent the data distribution, with white represented a pre-pandemic baseline, orange-red representing closures in UCLA and Los Angeles as whole, yellow representing the lifting of some but not all restrictions, and green representing full reopenings and the end of the anthropause. Lines within the violin plots represent 25%, 50% (thicker center line), and 75% quantiles. Data points per category are jittered. Groups with the same letter are not statistically significantly different from each other. Groups with different letters are statistically significantly different.
